# Novel therapy for gastric cancer peritoneal dissemination using genetically modified dental pulp cells and astatine-211

**DOI:** 10.1101/2025.10.15.682497

**Authors:** Komei Kuge, Wan-Ying Du, Hiroki Masuda, Tomohiko Yasuda, Toshifumi Tatsumi, Xiaojie Yin, Masao Yoshino, Akira Sugiyama, Ryohei Numata, Wataru Yokoyama, Hidenori Shibahara, Tomoko Tanaka, Mika Kobayashi, Akashi Taguchi, Takahide Kohro, Naoko Takubo, Masahiro Nakamura, Jaewoong Jang, Yoshitaka Kumakura, Hiroshi Yoshida, Akira Yoshikawa, Hiromitsu Haba, Youichiro Wada, Kiwamu Imagawa, Sachiyo Nomura

**Affiliations:** Department of Gastrointestinal Surgery, Graduate School of Medicine, The University of Tokyo; Department of Surgery, Graduate School of Medicine, Nippon University; Isotope Science Center, The University of Tokyo; Nishina Center for Accelerator-Based Science, RIKEN; Institute for Materials Research, Tohoku University; Clinical Pharmaceutical Sciences, School of Pharmacy and Pharmaceutical Sciences, Hoshi University; Regenerative Medicine Unit, Drug Discovery Research Institute, Research Division, JCR Pharmaceuticals Co., Ltd.; Department of Clinical Informatics, Jichi Medical School; Department of Diagnostic Radiology and Nuclear Medicine, Saitama Medical Center, Saitama Medical University; New Industry Creation Hatchery Center, Tohoku University

## Abstract

Peritoneal dissemination of gastric cancer is a terminal stage with limited treatment options and a five-year survival rate below 10%. To develop a more effective treatment, we created a new approach that uses genetically engineered human dental pulp cells (DPCs) that express the sodium/iodide symporter (NIS). These cells (NIS-DPCs) deliver astatine-211 (At-211) directly to tumor sites. When injected intraperitoneally into a mouse model of gastric cancer dissemination, DPCs, which were isolated and expanded, showed strong tumor-homing ability. Chemotaxis mediated by the CXCR4/SDF-1 axis has been reported to play a role in this accumulation. In addition to histological analysis, fluorescent imaging confirmed the selective accumulation of NIS-DPCs within tumor lesions. Introducing the NIS gene markedly increased SLC5A5 expression, enabling efficient At-211 uptake. To avoid nonspecific binding, sodium At-211 was utilized. High-resolution alpha-particle imaging visualized alpha-ray emission specifically from NIS-DPCs, confirming the radionuclide’s intracellular retention. *In vivo*, the sequential administration of NIS-DPCs, followed by Na[At-211], resulted in the pronounced regression of peritoneal tumors and a significant extension of survival compared to controls. The therapeutic mechanism involves three coordinated steps: tumor-directed migration of NIS-DPCs, At-211 uptake via NIS transporters, and localized alpha-particle–mediated cytotoxicity. This study introduces the novel concept of cell-based alpha-radiotherapy, which integrates regenerative and nuclear medicine. Due to the well-documented safety profile of DPCs in human clinical trials (J-REPAIR, jRCT1080224505), the NIS-DPC platform emerges as a promising approach for the precise, localized irradiation of disseminated gastric cancer and other challenging malignancies.

## INTRODUCTION

Peritoneal dissemination, a common terminal manifestation of gastric cancer, is classified as stage IV disease. It has a 5-year survival rate of less than 10%. Current therapeutic options are largely palliative, highlighting the urgent need for new treatment strategies with curative potential (Ref. Kanda, 2020).

Previously, we established a murine gastric cancer peritoneal dissemination cell line (YTN16) (Ref. Yamamoto 2018) and demonstrated the efficacy of targeted alpha-particle therapy using astatine-211 (At-211) (Ref. Watabe, 2025), a radionuclide with a high linear energy transfer and a 7.2-hour half-life. An anti-fibroblast growth factor receptor 4 (FGFR4) antibody, selected for its high expression on YTN16 cells, was conjugated to At-211 via a halogen-tin exchange reaction. *In vivo* biodistribution studies in a syngeneic C57BL/6J peritoneal dissemination model revealed that intraperitoneal administration resulted in peak peritoneal radioactivity at 4.5 hours. This treatment produced significant tumor regression and extended survival compared with untreated controls.

However, the therapeutic window is limited by the time required for antigen-antibody binding (approximately 24 hours) and the physical half-life of At-211. To improve delivery kinetics, we sought a vehicle capable of rapidly homing to the tumor microenvironment. Tumors actively recruit stromal fibroblasts to form cancer-associated fibroblasts (CAFs), and mesenchymal stem cells (MSCs) have similar tumor-tropic properties (Ref. Rosu, 2024). We investigated dental pulp cells (DPCs), developed by JCR Pharmaceuticals, as a potential vehicle. DPCs have an established manufacturing process and have been demonstrated to be safe in a phase 1/2 clinical trial for acute ischemic stroke (Ref. jRCT1080224505).

In this study, we evaluate NIS-DPCs as a biologic vector for targeted alpha-particle therapy in gastric cancer peritoneal dissemination. We hypothesize that leveraging the tumor-homing ability of NIS-DPCs can facilitate the delivery of At-211 to the tumor niche, thereby overcoming the limitations of radioimmunotherapy and enabling improved therapeutic efficacy and survival outcomes.

## RESULTS

### Preparation and characteristics of DPCs

Dental pulp cells (DPCs) are located in dental pulp, which is the soft connective tissue found inside teeth (Figure 1a). These cells play a key role in tooth vitality by producing dentin, supporting sensory functions, and participating in immune defense and tissue repair (Ref. Nito). Following dental pulp extraction, mild enzymatic digestion was performed to loosen the extracellular matrix. Then, mechanical mincing produced small tissue pieces. These explants were placed in culture to allow for the maintenance and expansion of DPCs (Ref. Gronthos) (Figure 1b). DPCs were able to expanded using a culture medium; however, no proliferation was observed in soft agar, suggesting that they were not tumorigenic when injected *in vivo* (Figure 1c).

**Figure 1.**
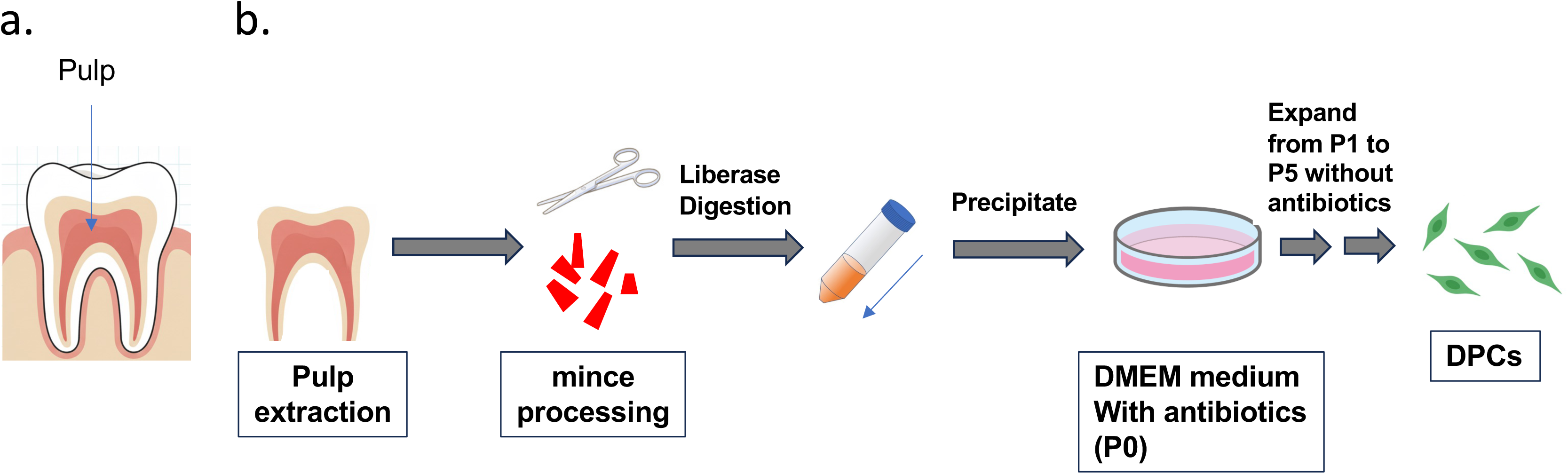
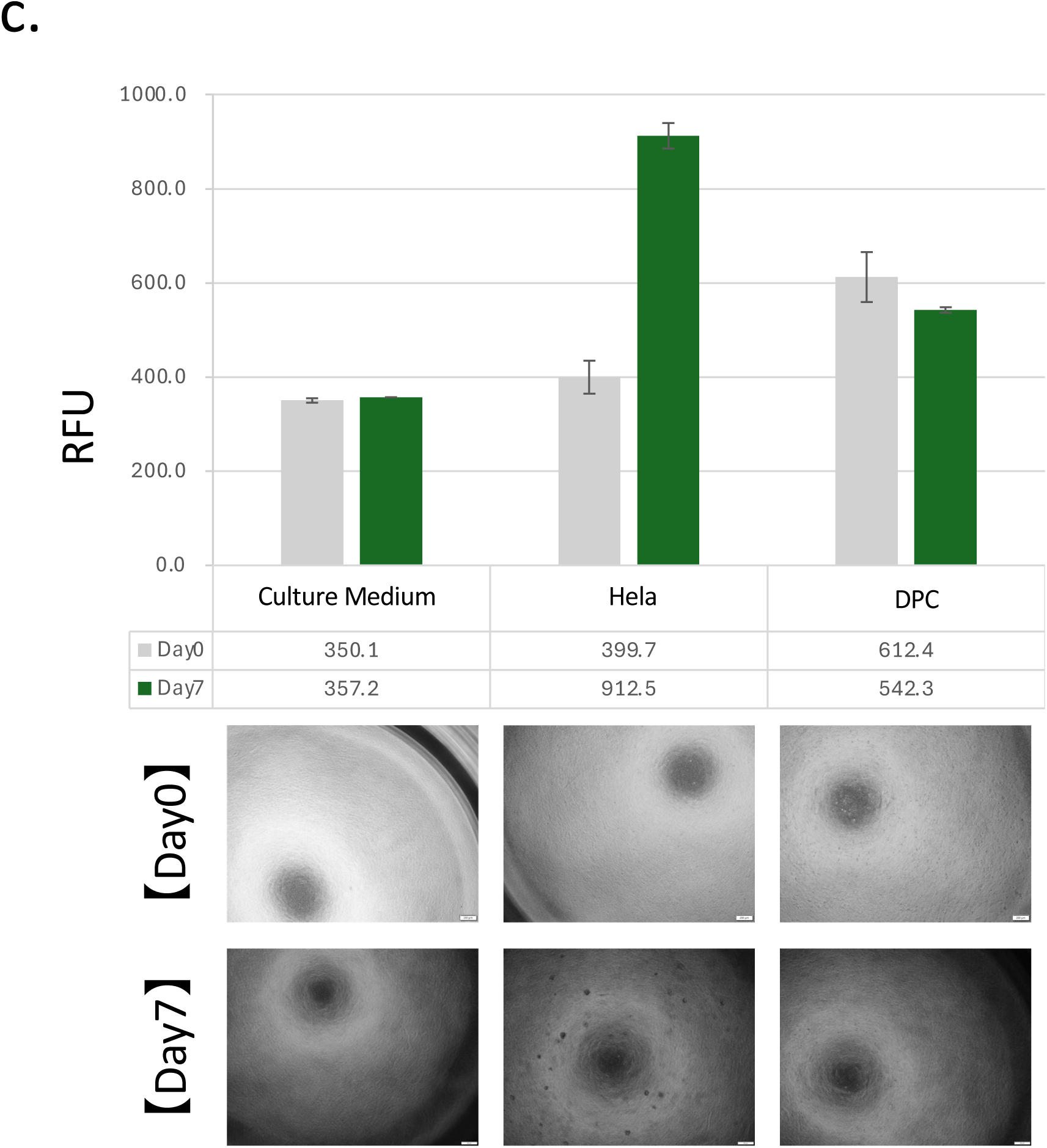
Isolation and characterization of dental pulp cells (DPCs). (a) The dental pulp is a soft connective tissue located within the interior of the tooth. (b) After extraction of the dental pulp tissue, the connective components were enzymatically digested, mechanically minced with a sharp blade, and cultured in growth medium. Proliferating cells were isolated and purified as DPCs. (c) Colony-forming assay of DPCs. HeLa cells were used as a control. Cells were cultured for 0 or 7 days, then stained with a fluorescent dye. Fluorescence intensity was quantified as relative fluorescence units (RFU; n = 3). Representative microscopic images of colonies are shown. Scale bar, 200 μm.

### Confirmation of DPC accumulation in peritoneal tumor tissue

To evaluate the specificity of tumor-directed accumulation, 1.0 × 10⁶ DPCs were injected intraperitoneally into a previously established gastric cancer mouse model (Ref. Nagaoka). Two days after the final DPC injection, the mice were sacrificed and their peritoneal tissues were fixed and stained immunohistochemically using an anti-human Lamin B1 antibody. Brown staining, which indicates DPC localization, was observed in tissues surrounding established tumor lesions (Figures 2a and 2b), whereas no staining was detected in tissues lacking tumor cell engraftment (Figure 2c). These results suggested that intraperitoneally administered DPCs migrate to and accumulate within peritoneal dissemination sites.

**Figure 2.**
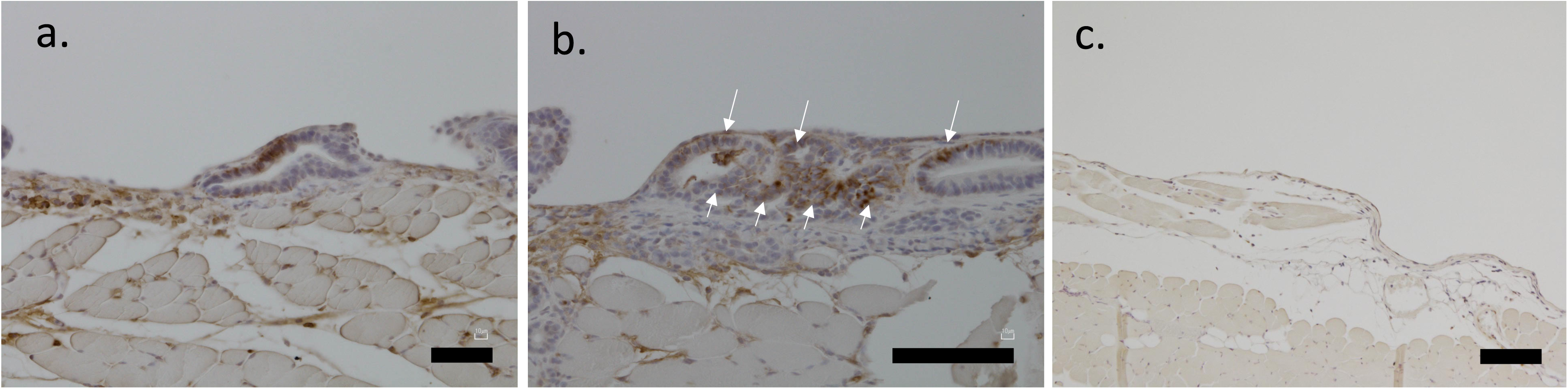
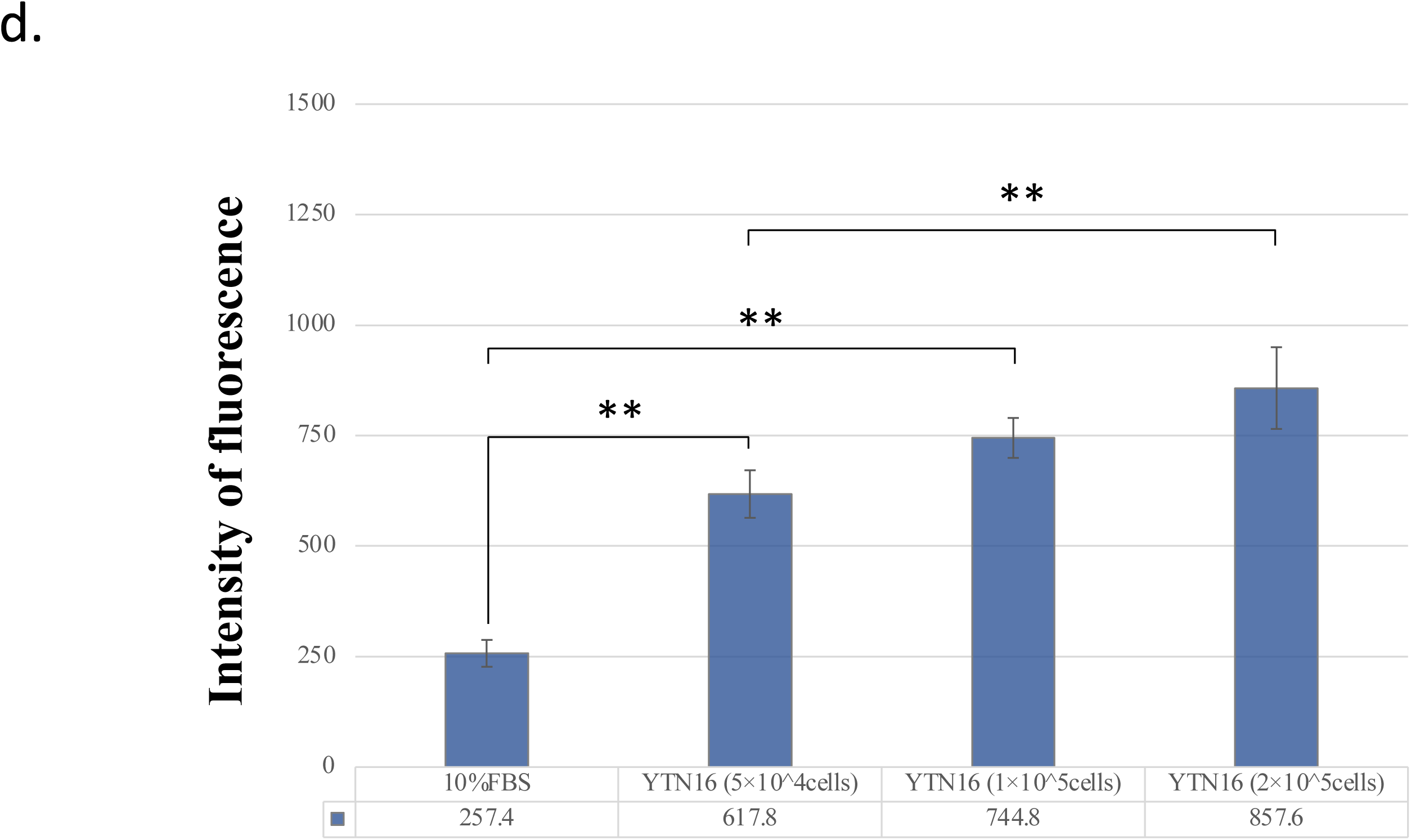
Confirmation of DPC accumulation in peritoneal tumor tissue. (a–c) Histological observation of peritoneal tumor tissue 1.5 weeks after intraperitoneal administration of DPCs. Nuclei were counterstained with hematoxylin and eosin stain, and DPCs were detected by immunostaining using an anti-human Lamin B1 antibody followed by an HRP-conjugated secondary antibody. Panel (a) shows a low-magnification image, and (b) shows a high-magnification view of the region. Arrows indicate DPCs accumulated within peritoneal tumor lesions. Panel (c) shows tissue from a normal mouse injected with DPCs as a control. (scale bars=50 µm in a&c, scale bar=100 µm in b.) (d) Migration assay evaluating DPC chemotaxis toward YTN16 gastric cancer cells. The lower chamber contained 5 × 10⁴ to 2 × 10⁵ YTN16 cells, and 1 × 10⁵ DPCs were seeded in the upper chamber. After 48 hours, calcein staining was performed and fluorescence intensity of migrated cells was quantified. DPC migration increased in proportion to the number of YTN16 cells in the lower chamber (n = 4; **p < 0.01).

To validate the migratory ability of DPCs toward the YTN16 cell line, a migration assay was performed using a Transwell chamber. After 1.0 × 10⁵ DPCs were placed in the upper chamber, different numbers of YTN16 cells were placed in the lower chamber. The cells that migrated to the other side of the micropore membrane were stained with a calcein solution, and the absorbance was measured. Figure 2d shows that the presence of YTN16 cells induces DPC migration in a cell number-dependent manner, proving the migratory capability of DPCs towards YTN16 cells.

### Functional transduction of NIS to DPC

The sodium-iodide symporter (NIS) is a transmembrane glycoprotein with 13 transmembrane domains that transports radioactive isotopes, such as I-131, I-123, I-125, I-124, Tc-99m, Re-188, and At-211, into cells (see reference by Darrouzet). We introduced the NIS gene (SLC5A5) into DPC cells using lentivirus. Although the expression levels of other NIS-encoding genes, such as SLC26A4 and SLC26A7, remained unchanged, the expression level of SLC5A5 in NIS-DPC increased by over 100-fold (Figure 3a).

**Figure 3.**
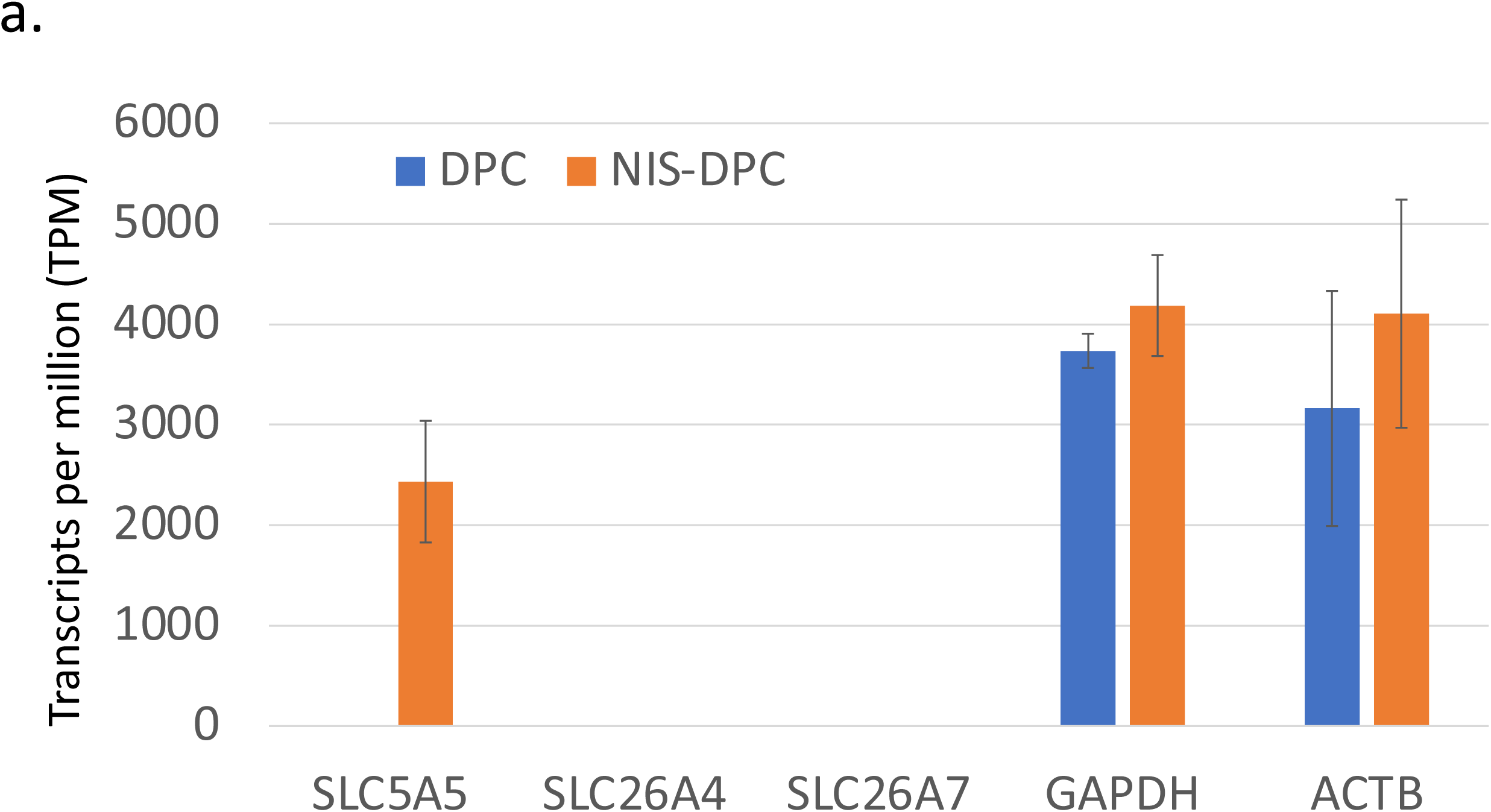
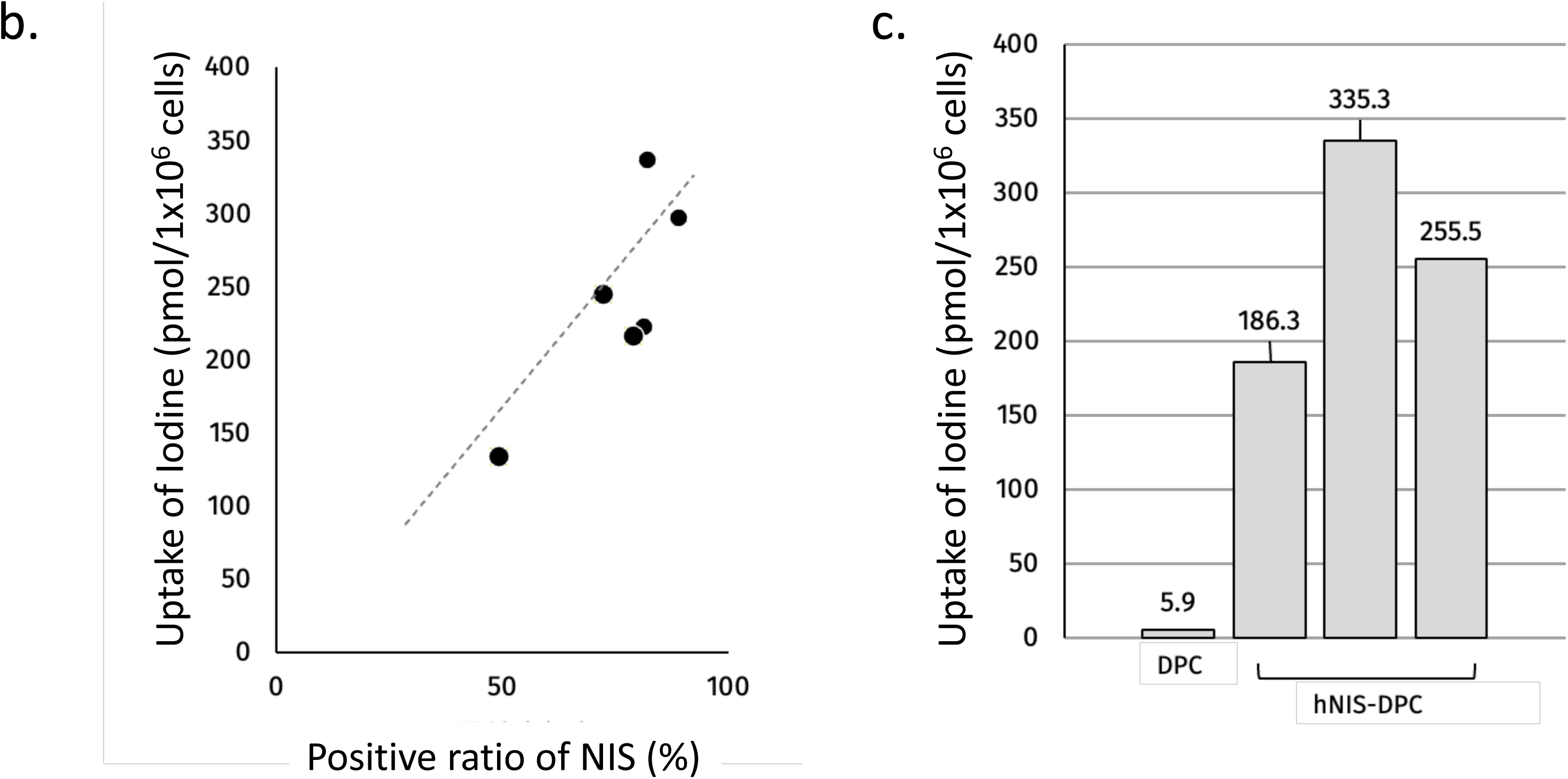
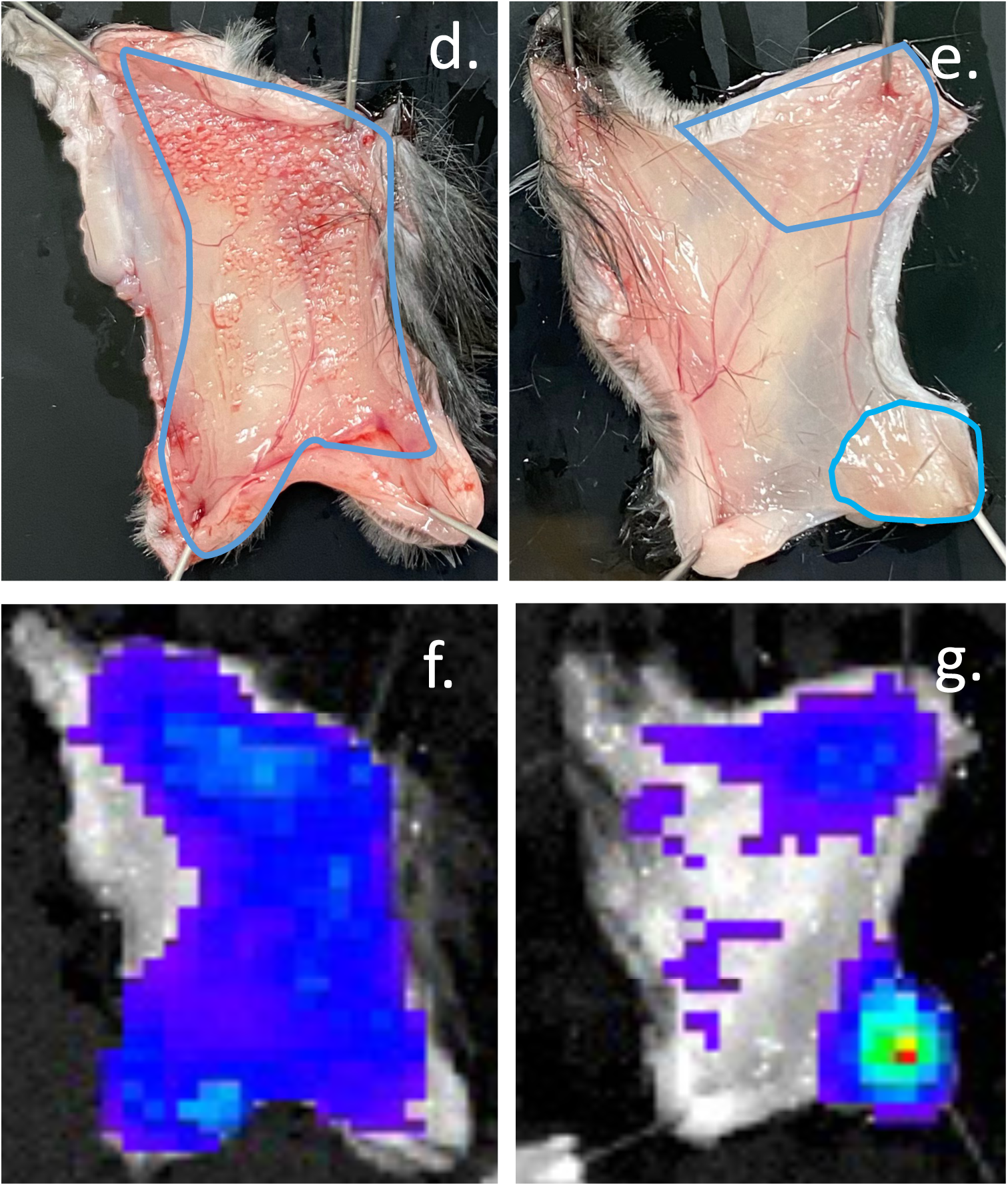
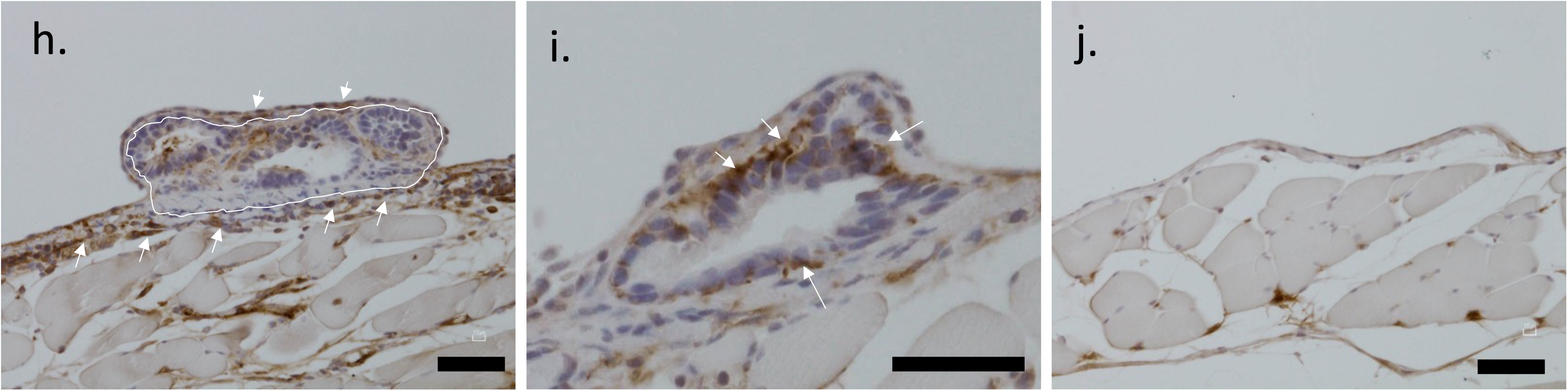
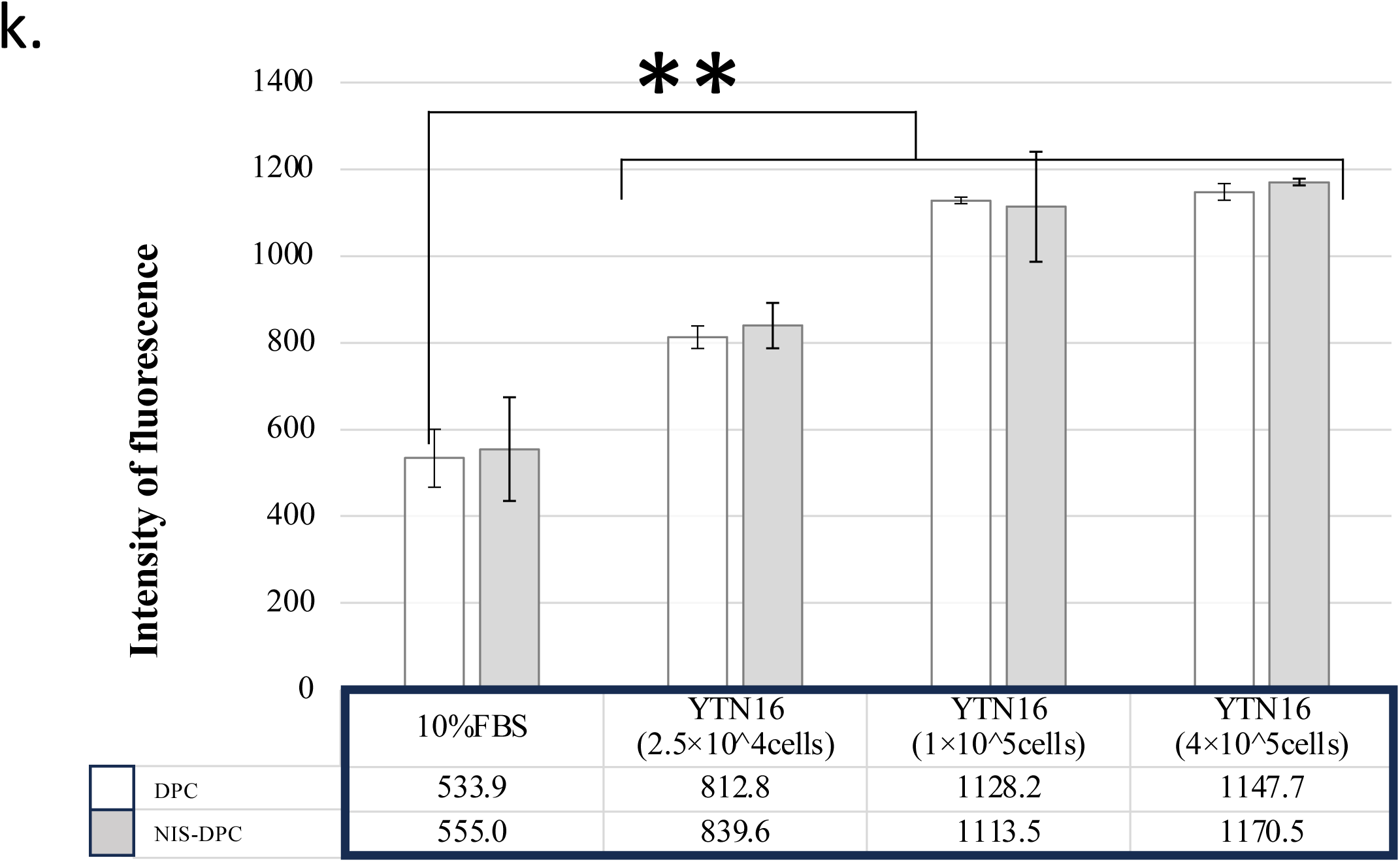
Functional transduction of the sodium/iodide symporter (NIS) into DPCs. (a) Gene expression levels of parental DPCs and sodium/iodide symporter (NIS)–transduced DPCs (NIS-DPCs) quantified from RNA-seq data and presented as transcripts per million (TPM) values. Among iodide symporter genes, SLC5A5 was transduced and expressed in NIS-DPC. GAPDH and beta-actin (ACTB) were controls (n=3). (b) Correlation analysis between NIS-positive cell fraction and iodide uptake levels in NIS-DPCs. (c) Comparison of iodide uptake between DPCs and NIS-DPCs. Data for NIS-DPCs were obtained from different lots. (d–g) Macroscopic and fluorescence imaging analyses of a gastric cancer peritoneal dissemination model after intraperitoneal administration of DiR-labeled DPCs. Panels (d) and (e) show macroscopic views of the peritoneal serosa, where blue lines outline the regions of peritoneal tumor nodules. Corresponding in vivo imaging system (IVIS) fluorescence images (f and g) demonstrate colocalization of fluorescence signals with the tumor lesions identified in (d) and (e). (h∼j) *In vivo* localization of NIS-DPCs within tumor tissue, showing accumulation comparable to that of unmodified DPCs. Scale bars=50 µm in h&j, scale bar=100 µm in i. (k) Migration assay using Transwell chambers with YTN16 gastric cancer cells placed in the lower compartment. NIS-DPCs exhibited cell number–dependent migration similar to DPCs (n = 3; **p < 0.01).

Next, we verified the expression level of NIS molecules on the cell surface. Then, we evaluated the correlation between uptake capacity and positive rate using iodine as a surrogate molecule *in vitro*. The results are shown in Figures 3b and 3c. We observed that iodine uptake increased in a manner dependent on the NIS-positive rate and observed a trend in which higher NIS-positive rates correlated with higher iodine uptake (Figure 3b). When we measured iodine uptake in three lots of NIS-DPC cells, we observed more iodine uptake was observable compared to DPC cells without gene introduction. These results confirmed that the introduced NIS gene was expressed on the cell surface and that its function was maintained (Figure 3c).

To investigate whether NIS-DPCs could preserve the ability to accumulate in tumors, we first compared the localization of disseminated lesions and NIS-DPCs using fluorescent dye staining. Briefly, 5.0 x 10⁵ cells of NIS-DPCs were prepared as described in the Materials and Methods section. Then, the cells were labeled with a fluorescent dye (DiR) for live cells, and were administrated into peritoneal cavity of dissemination models. After 24 hours, the mouse models were euthanized and their peritoneal surfaces were opened to apply a fluorescence detection camera. The localization of the injected NIS-DPCs was visualized using an *in vivo* imaging system (IVIS Lumina III) and was compared with the distribution of disseminated visible lesions. As shown in Figures 3d–3g, the area occupied by the lesion was surrounded by a blue line (left in Figure 3d and right in Figure 3e). Fluorescent images obtained using a fluorescent camera are shown in Figures 3f (left) and 3g (right). Significant colocalization was observable on both sides, suggesting the accumulation of NIS-DPCs in gastric cancer cells. To validate the ability of NIS-DPCs to migrate into dissemination lesions, NIS-DPCs were injected into the peritoneal cavity in a manner. Two days after injection, the peritoneal tissues were subjected to immunohistochemical staining using an anti-human lamin B1 antibody. Brown staining, which indicates NIS-DPC localization, was observed in tissues surrounding established tumor lesions (Figures 3h and 3i), whereas no staining was detected in tissues lacking tumor cell engraftment (Figure 3j). A migration assay revealed significant, cell-number-dependent migration of NIS-DPCs to YTN16 cells, similar to what was observed with DPCs (Figure 3k). These findings strongly suggest that NIS-DPCs administered intraperitoneally accumulate within peritoneal dissemination lesions.

### Uptake of At-211 by NIS-DPC

Recently, it was reported that purified At-211 undergoes nonspecific binding *in vivo* due to its various chemical forms, and that this can be controlled by converting it to sodium At-211 (Ref. Watabe, 2022). Therefore, prior to *in vivo* administration, we prepared sodium At -211 using the method shown in Figure 4a. We confirmed the uniform chemical form of the purified At-211 using thin-layer chromatography (TLC) (Figure 4b).

**Figure 4.**
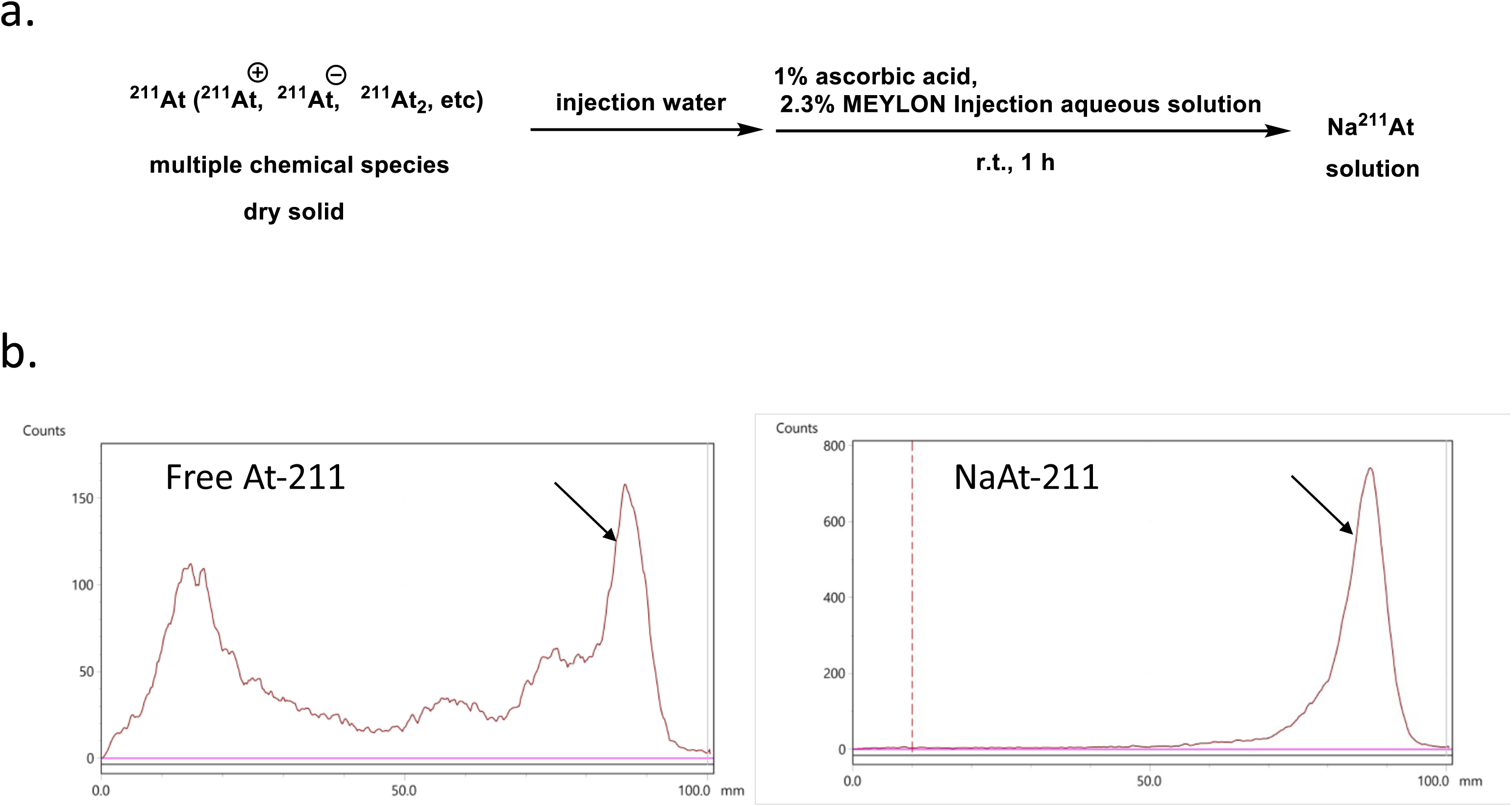
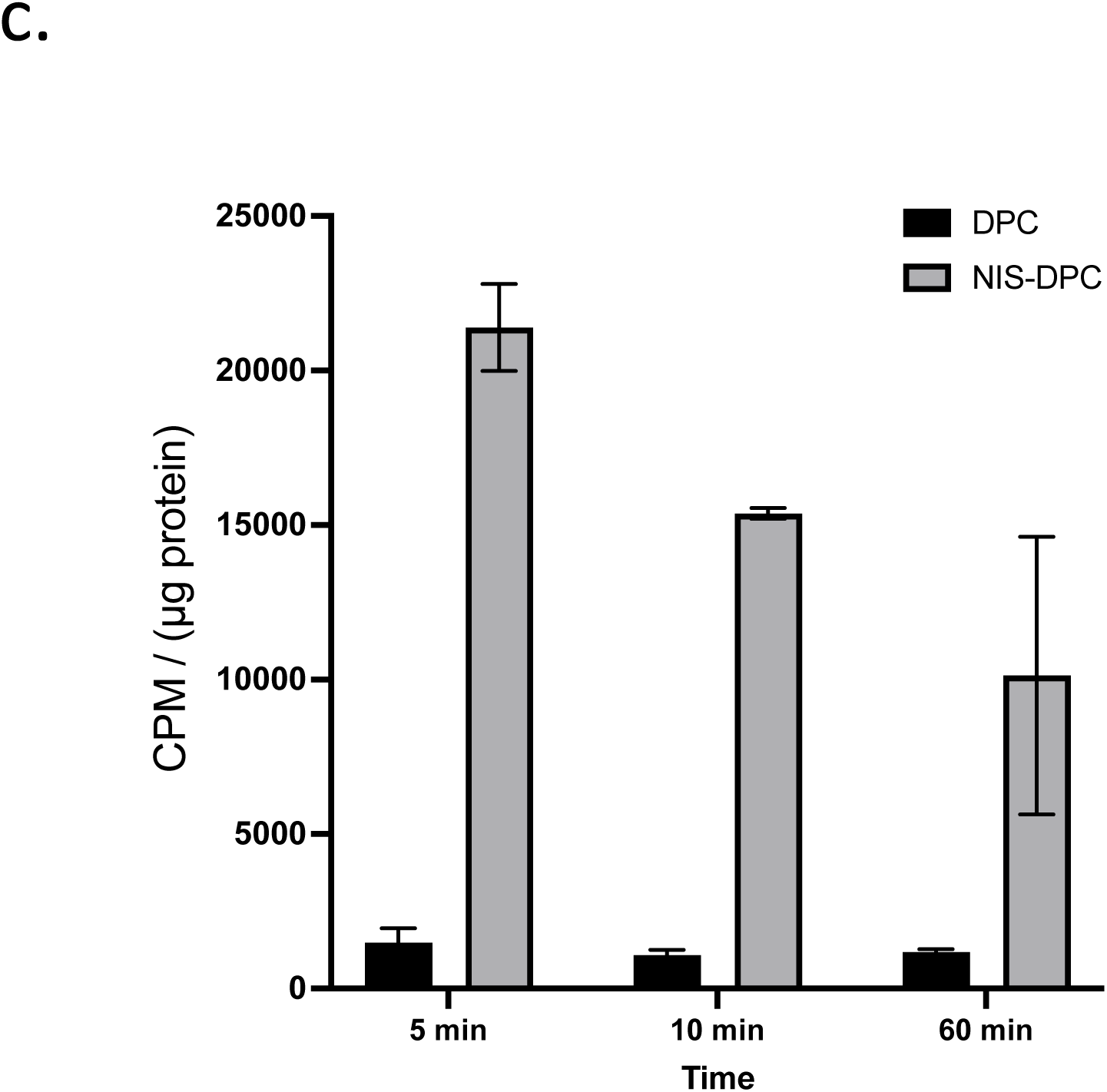
Preparation and functional characterization of sodium astatide-211 (Na[At-211]) and its uptake by NIS-DPCs. (a) Reaction protocol for converting purified free At-211, produced at an accelerator facility, into a homogeneous sodium astatide form (Na[At-211]). The free isotope can exist in multiple chemical species, and sodium conjugation was performed to stabilize its form. (b) Thin-layer chromatography (TLC) analysis of At-211 before and after sodium conjugation. Left: TLC profile of free At-211 (∼100 kBq). Right: TLC profile of Na[At-211] (∼280 kBq). Arrows indicate the Na[At-211] peak position. (c) Cellular uptake of Na[At-211] by DPCs and NIS-DPCs. DPCs or NIS-DPCs (5 × 10⁴ cells per well) were seeded in 24-well plates and incubated with 10 kBq Na[At-211]. At 5, 10, and 60 min, cells were collected, fractionated, and measured for protein concentration and radioactivity. Radioactivity normalized to protein content was calculated. NIS-DPCs exhibited approximately tenfold higher At-211 uptake than parental DPCs (n = 3; black bars, DPCs; gray bars, NIS-DPCs).

Furthermore, we determined the amount of At-211 taken up by DPCs and NIS-DPCs. The results are shown in Figure 4c. Similar to the results of confirming uptake capacity using a surrogate molecule, NIS-DPC cells with the introduced NIS gene took up more At-211 than DPC cells without the gene. Additionally, the highest uptake was observed five minutes after adding At-211 when comparing changes in uptake amount over time at five, ten, and sixty minutes. These results suggest that At-211 is rapidly taken up by NIS-DPC cells via the NIS protein. Therefore, pre-accumulating NIS-DPC cells around the peritoneal dissemination site and then administering At-211 enables the alpha-emitters to be quickly taken up by the cells and be effective within their half-life.

### Novel contact scintillator imaging of alpha-ray trajectories from alpha emitters

Because the NIS-DPC cell fraction was radioactive, uptake of At-211 was suggested in Figure 4c. However, to verify that the alpha-rays emitted from the At-211 originate from the NIS-DPC cells, a camera capable of visualizing alpha-rays on a 1-μm scale was used. Figure 5a shows the alpha camera system configuration (Refs. Yamamoto, 2023; Yoshino, 2025). The system comprises three principal components: (i) a 100-μm-thick GAGG thin-film scintillator screen, (ii) an infinity-corrected magnification optical train consisting of an objective and an imaging lens, and (iii) a CMOS imaging sensor (specifically the ORCA-Quest 2 qCMOS [C15550-22UP] from Hamamatsu Photonics). Alpha-particles emitted from within the cells are converted into photons by the GAGG scintillator, magnified by the 20x objective lens, and detected by the CMOS sensor. Figure 5b shows an overview of the system. Eighteen thousand NIS-DPC and DPC cells were cultured in an eight-well chamber slide. Then, 500 kBq of At-211 was added to the culture medium per well, followed by a 30-minute incubation period. The cells were fixed with formalin and observed with an alpha camera for ten minutes. Figures 5c and 5f show overviews of the NIS-DPC and DPC cells, respectively. Both exhibit similar morphology and distribution. However, Figures 5d and 5g show that the alpha-particle range was detected only in the NIS-DPC cells. Figures 5e and 5h are merged images of cells and alpha-rays.

**Figure 5.**
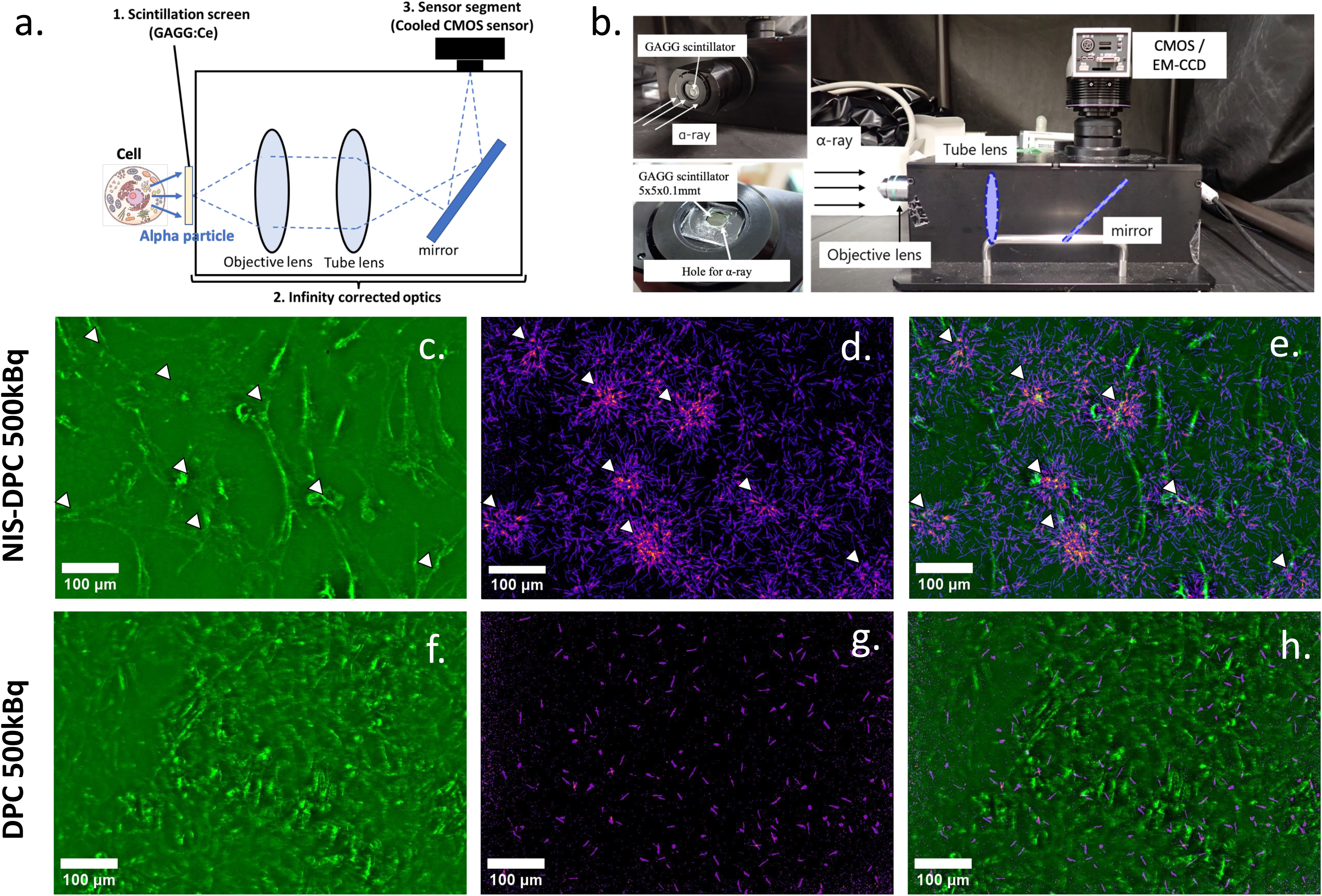
Detection of alpha-particle emission from NIS-DPCs. (a) Schematic diagram of the alpha-particle detection system. Alpha-particles emitted from cells (left) are first converted into photons by a GAGG:Ce scintillator. The photons are then amplified and reflected within the detector and captured by a cooled CMOS sensor positioned at the top. (b) Overview of the alpha-particle detection device. (c–e) NIS-DPCs (1.8 × 10⁴ cells per well) were seeded in 8-well chamber slides and incubated for 30 min with 500 kBq of Na[At-211]. Cells were then fixed with formalin and imaged. Arrowheads indicate the localization of individual cells. (f–h) DPCs (1.8 × 10⁴ cells per well) were similarly treated with 500 kBq Na[At-211], incubated for 30 min, fixed with formalin, and imaged. (c and f) Optical microscopic images of cells. (d and g) Corresponding alpha-camera signal images. (e and h) Merged images combining optical and alpha-particle signals. Scale bar=100 μm.

Colocalization of alpha-ray and cell images was only observable in NIS-DPC cells treated with At-211. These results suggest that the NIS-DPC cells efficiently took up At-211 and emitted alpha-rays.

### At-211 and NIS-DPC cell administration showed antitumor effects

Then, we evaluated the antitumor effects and survival prolongation associated with the combined administration of NIS-DPCs and At-211 using a mouse peritoneal dissemination model. Two weeks after tumor cell transplantation, we administered 0.6 MBq of At-211 per animal. Twenty-four hours prior to the At-211 injection, 0.5 × 10⁶ NIS-DPC cells were administered intraperitoneally. One and a half weeks after transplantation, examination of the peritoneum revealed tumor cells in the peritoneal dissemination group (Figure 6a), the NIS-DPCs administration group (Figure 6b), and the DPCs and At-211 administration group (Figure 6c). However, disappearance of peritoneal seeding was observed in the group that received both NIS-DPCs and At-211 (Figure 6d, asterisks).

**Figure 6.**
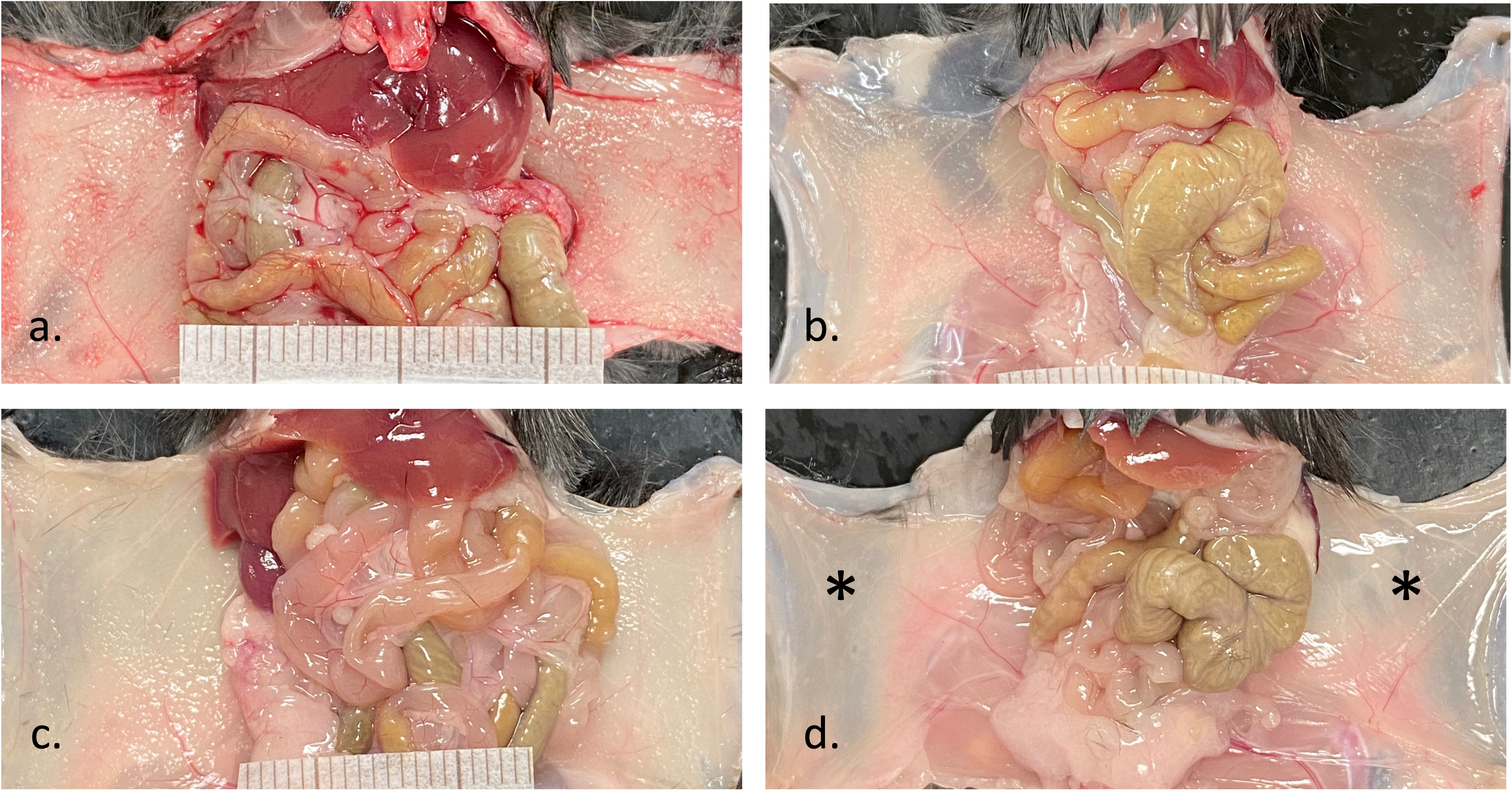
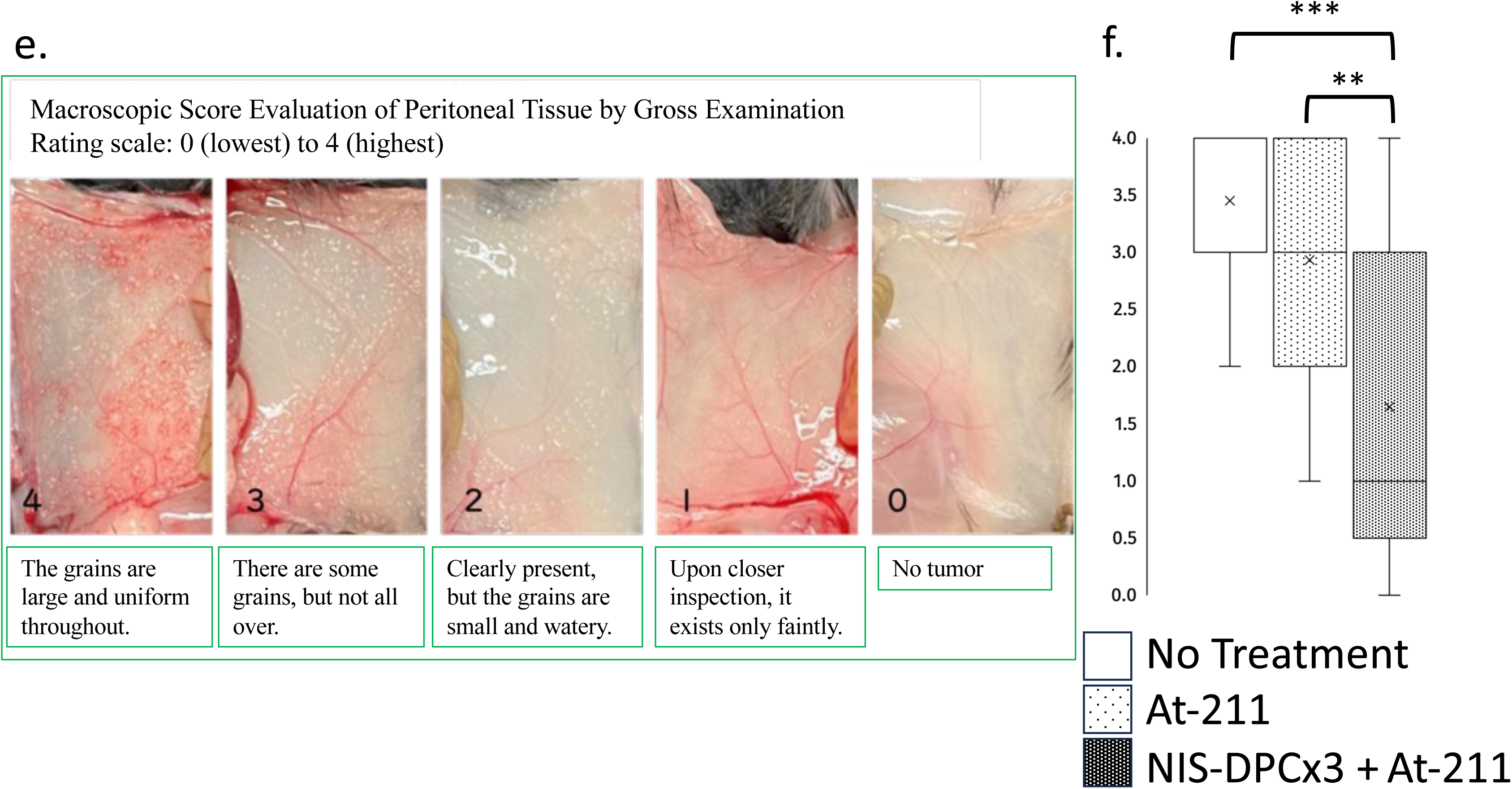
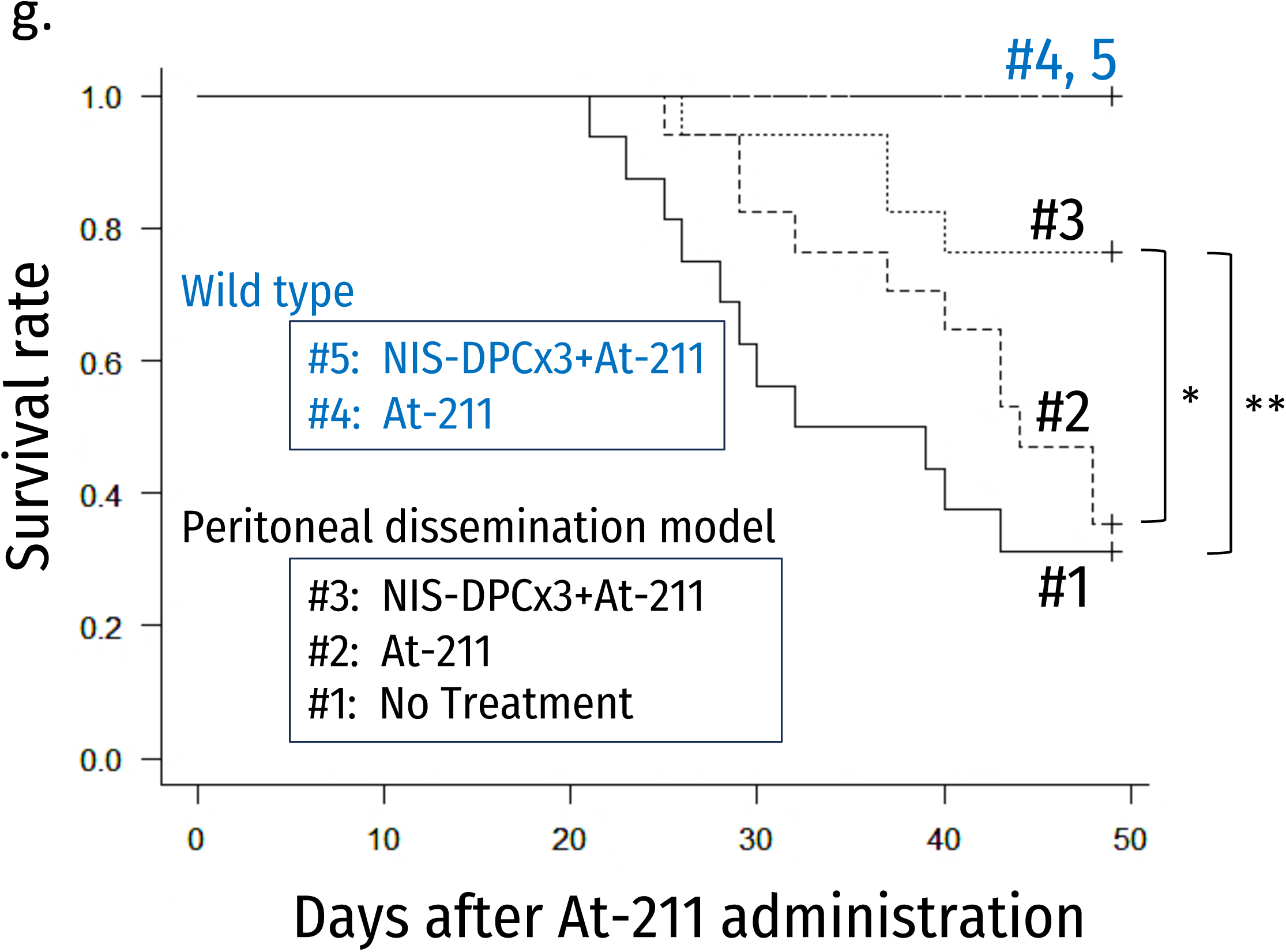
Therapeutic efficacy of NIS-DPCs combined with At-211 in a murine gastric cancer peritoneal dissemination model. To establish a peritoneal dissemination model, C57BL/6 mice were inoculated intraperitoneally with 1 × 10⁷ YTN16 gastric cancer cells. Two days prior to nucleotide injection, 0.5 × 10⁶ NIS-DPCs were administered intraperitoneally. Then, two weeks after YTN16 transplantation, 0.6 MBq of Na[At-211] was injected intraperitoneally. The mice were sacrificed 1.5 weeks later for macroscopic observation. (a) Untreated control mouse. (b) A mouse that received an intraperitoneal injection of NIS-DPCs only. (c) A mouse that was treated with DPCs and Na[At-211]. (d) A mouse that received an intraperitoneal administration of NIS-DPCs, followed by Na[At-211]. The peritoneal tumor burden disappeared after treatment with NIS-DPCs and Na[At-211]. Asterisks indicate a smooth peritoneum. (e) Macroscopic scoring of peritoneal dissemination was performed on a 0–4 scale. (f) Quantitative comparison of macroscopic scores among the untreated group, Na[At-211]-only group, and the group receiving three NIS-DPC administrations plus Na[At-211]. Statistical analysis was performed using Tukey’s test (n = 10; **p < 0.01, ***p < 0.001). (g) Kaplan–Meier survival analysis after one month following Na[At-211] administration. Mice treated with three administrations of NIS-DPCs plus Na[At-211] (#3) showed a significant survival advantage compared with the untreated group (#1) and the Na[At-211]-only group (#2). Control groups without YTN16 cell implantation included Na[At-211]-only (#4) and NIS-DPC × 3 + Na[At-211] (#5) treatments (#1 : n = 16, #2/3 : n = 17, #4/5 : n = 3; *p < 0.05, **p < 0.01).

We evaluated the effect of NIS-DPC & At-211 treatment by scoring the antitumor effects visually on a 5-point scale (0–4). This scoring was based on the extent of cancer cells adhering to the peritoneal tissue at the time of death or at the end of the observation period (Figure 6e).

Using the same evaluation scale, we performed a series of experiments to optimize the At-211 radioactivity, NIS-DPC dose, and NIS-DPC injection repetition (data not shown). Based on macroscopic evaluation of the extent of peritoneal dissemination, a significantly higher score was achieved with 0.6 MBq of At-211/body, 5x10⁵/body of NIS-DPCs, and three administrations of NIS-DPCs (data not shown).

The results of the antitumor effect of the optimized procedure are analyzed using Turkey test and are shown in Figure 6f as a box-and-whisker plot. Compared to the untreated group (no treatment) and the At-211 monotherapy group (At-211), a significant reduction in score (i.e., reduction in disseminated tissue) was observed in the NIS-DPCx3+At-211 group (NIS-DPCx3+At-211). Therefore, it was suggested that the administered NIS-DPC accumulated around the implanted cancer cells in the peritoneum. Subsequently, the cancer tissue was reduced by the cytolytic effect of alpha-rays after uptake of the administered At-211.

We evaluated the survival rates using this optimized regimen with Kaplan-Meier curves and Logrank tests. The survival rates of individual mice in each group were evaluated after At-211 administration. The results are shown in Figure 6g. Compared to the untreated and At-211-treated groups, the NIS-DPCx3+At-211 group showed significantly longer survival.

These results suggest that the extension of the survival rate is due to the reduction and damage of the implanted tumors by NIS-DPCx3+At-211 in the mouse peritoneal dissemination model.

## DISCUSSION

In this study, we created NIS-DPC cells by forcing the expression of the NIS gene— which encodes an iodine transporter—in DPC cells that accumulate in the peritoneal cavity. Forty-eight hours after administering the cells, we introduced At-211 into the peritoneal cavity. This demonstrated that the NIS-DPC cells accumulated around the tumor cells rapidly took up At-211, resulting in tumor cell death. Figure 7 illustrates this three-step treatment method: addition of NIS-DPC cells, addition of At-211, and alpha-ray irradiation. The first step involves accumulating NIS-DPC cells in tumors. The second process involves At-211 uptake by NIS-DPC cells. The third process involves tumor-killing effects.

**Figure 7.**
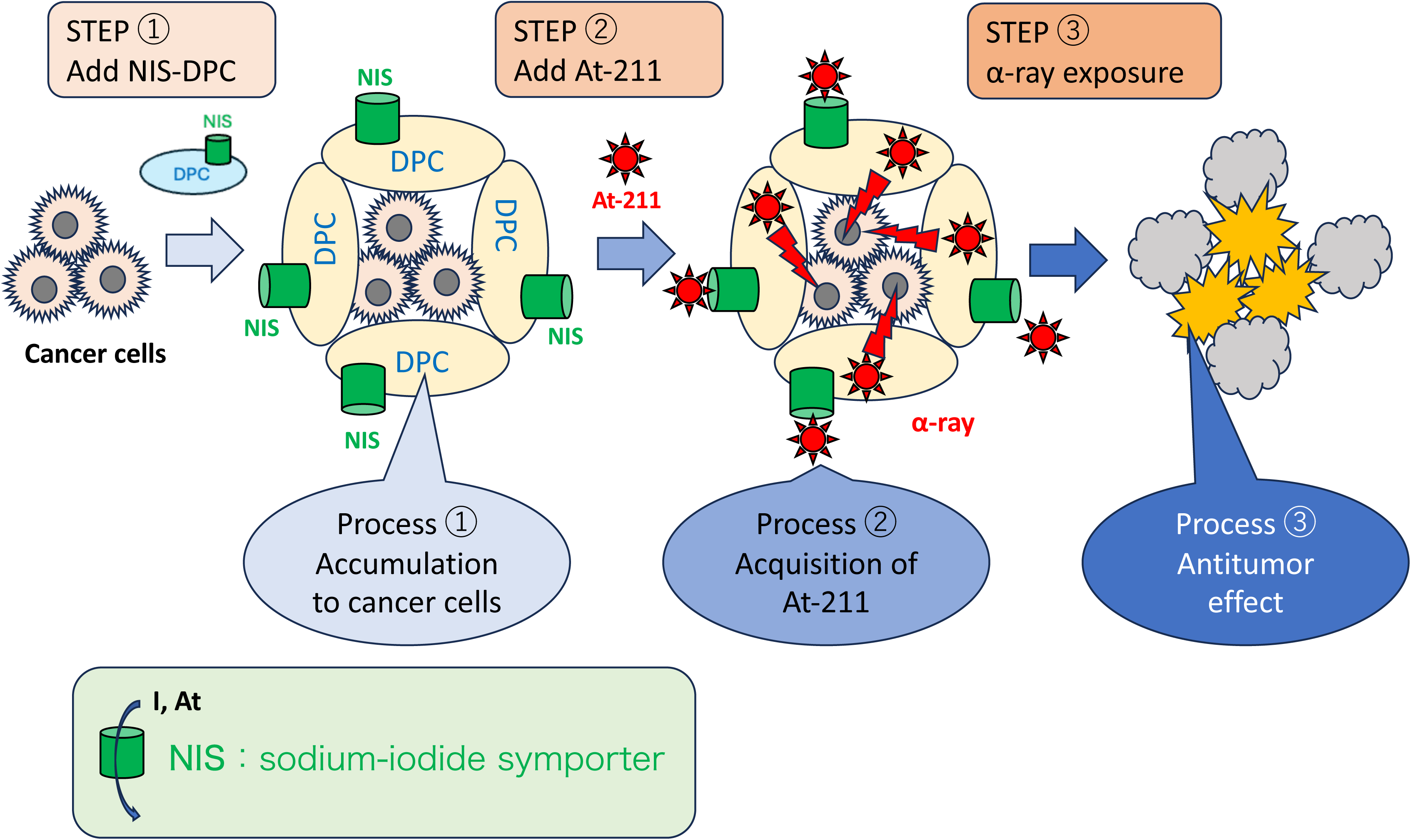
Graphical Summary. Stepwise overview of the NIS-DPC cancer-targeting concept. The schematic illustrates the translational workflow of NIS-DPCs as a targeted theranostic platform for cancer therapy. Step 1 (engineering). DPCs were genetically modified to express the sodium/iodide symporter (NIS), which mediates the cellular uptake of both iodide and astatine. The lower panel depicts this dual transport function of NIS, highlighting its ability to internalize iodide and the αalpha-emitting radionuclide At-211. Step 2 (targeting and accumulation). Following administration, NIS-DPCs preferentially accumulated in cancer cells, enabling the selective delivery of At-211 to tumor sites. Step 3 (localized therapy/outcome). Localized α radiation at the tumor site induced targeted cytotoxicity, demonstrating the translational potential of NIS-DPC–mediated therapy for precise cancer treatment.

### Accumulation of DPCs/NIS-DPCs to cancer cells

It is well-known that cancer cells accumulate large numbers of cancer-associated fibroblasts (CAFs) in their vicinity (Ref. Sahai, 2020). Furthermore, mesenchymal stem cells (MSCs) exhibit dynamics similar to those of CAFs (Ref. Antoon, 2024). MSCs migrate in response to chemokines produced by cancer cells, such as SDF-1 (Ref. Stoicov, 2013). MSCs recognize these chemokines via CXCR4. Due to protein cross-reactivity between humans and mice, however, human MSCs can also migrate toward mouse tumors. Additionally, MSCs have low immunogenicity and can be administered allogeneically. Since 2008, when it was reported that DPC cells (a type of MSC) express CXCR4 and migrate in response to SDF-1 (Ref. Jiang, 2008), numerous studies have reported that DPC cells accumulate in inflammatory or ischemic sites, particularly in oral and maxillofacial surgery (Refs. Yamaguchi, 2015; Li, 2017; Neves, 2020; Kornsuthisopon, 2022; Irfan, 2023). DPCs produced by JCR Pharmaceuticals have been administered to humans and proven safe (Ref. Suda, 2022). They were examined for their ability to migrate and accumulate around tumor tissues in a mouse peritoneal dissemination model (Ref. Nagaoka, 2025).

As mentioned above, the first step in this therapy is the accumulation of NIS-DPC cells in cancer cells. Multimodal experiments were performed to prove this. Macroscopically, as shown in Figures 3d to 3g, the localization of the fluorescently labeled NIS-DPC cells coincided with the localization of the gastric cancer peritoneal dissemination lesions. Microscopically, Figures 2a-2c and 3h-3j show that both DPCs and NIS-DPCs were present in gastric cancer lesions via immunohistochemical staining. These findings suggest that DPC/NIS-DPC cells accumulate in tumor cells.

*In vitro*, DPC/NIS-DPC cells were observed migrating toward gastric cancer cells (see Figures 2d and 3k). The molecular mechanism underlying this migration is thought to involve the induction of EGR1 in DPC/NIS-DPC cells by soluble factors released from the cancer cells (Ref. Xiao, 2021). Further clarification of this mechanism is needed in future studies.

### Functional uptake of At-211 by NIS-DPC

The second step of this treatment method involves NIS-DPC cells taking up At-211. Several experiments were conducted to determine whether the introduced iodine transporter functions in DPC cells. First, non-radioactive iodine and a non-radioactive ligand were added to DPC cells expressing NIS, and the uptake efficiency was examined. As shown in Figures 3, the amount of uptake increased in a NIS expression-dependent manner, demonstrating that NIS functionally takes up the ligand (Figure 3b), and NIS-DPCs exhibited approximately 50-fold higher iodine uptake than DPCs (Figure 3c). Furthermore, Figure 4c shows the efficient uptake of At-211 by NIS-DPCs beginning five minutes after At-211 administration. Using the newly developed alpha camera, we confirmed consistent uptake of At-211 and cell localization (Figure 5). This indicates that NIS-DPC cells take At-211 into the cell and emit alpha rays. In the future, we expect to directly visualize the lethal effect of alpha-rays on tumor cells by co-culturing NIS-DPC cells with tumor cells and observing them using the alpha camera.

### Tumoricidal effect of NIS-DPC&At-211

As shown in Figure 6, this study demonstrated that administering At-211 after three times NIS-DPC administration reduced peritoneal dissemination lesions and improved survival rate. This confirms the tumor-killing effect, the third process of this treatment. When evaluating the efficacy of anticancer drugs in humans, the primary endpoint is an extension of the survival period. Therefore, improvement in survival rate is equally important when evaluating the efficacy of cancer treatments using model animals as it is in humans. Furthermore, the proposed mechanism of action was strongly implicated. However, it has been reported that the immune response induced by neoantigens released from tumor cells damaged by radiation exposure contributes to the efficacy of nuclear medicine therapy (Ref. Nagaoka, 2025). Elucidating the mechanism in greater detail in the future may expand the application of combined therapy with NIS-DPC cells and At-211.

### Requirement for practical application

In order to commercialize this treatment method in the future, data on its efficacy must first be accumulated. Additionally, data on the safety and pharmacokinetics of the reagents used in this study will be necessary. Since DPC cells have previously been administered to humans and their safety has been demonstrated (Suda, 2022), data on the safety of NIS-DPCs is essential. In particular, since lentiviruses are used to introduce the NIS gene, it is necessary to determine if the viruses are excreted by the treated animals. Furthermore, since At-211 is administered as sodium astatine into the abdominal cavity, pharmacokinetic data must be obtained from medium-sized animals before human administration. Human clinical studies will be conducted based on these data.

DPC cells can be administered allogenically. Furthermore, they can be procured minimally invasively, and the cells can be stored frozen, making it feasible to establish a supply system. Meeting clinical demand for At-211 will be possible through the future widespread adoption of mass production technology and the development of dedicated accelerators.

### Perspective

Here, we reported a new treatment method combining nuclear and regenerative medicine products. Since the safety of DPC cells has already been proven in humans, a first-in-human trial should be conducted after the safety of NIS-DPCs and At-211 is evaluated.

## MATERIALS AND METHODS

### Preparation of Dental Pulp Cells (DPCs)

Human dental pulp cells (DPCs) were isolated from extracted teeth under sterile conditions, as previously described with minor modifications. Briefly, extracted teeth were first disinfected by immersion in 1% chlorhexidine gluconate solution followed by washing with sterile physiological saline to remove debris and surface contaminants.

The tooth structure was mechanically fractured using a dental rongeur, and the pulp tissue was carefully extracted from the pulp chamber. The obtained pulp tissue was minced into small fragments using sterile surgical scissors and transferred into a tube containing Liberase digestion solution (Roche Diagnostics, Basel, Switzerland). The tissue suspension was incubated at 37°C for enzymatic digestion with gentle agitation until the connective matrix was adequately loosened.

After digestion, the suspension was mixed with Dulbecco’s Modified Eagle Medium (DMEM) supplemented with 20% fetal bovine serum (FBS) and 1% penicillin/streptomycin (P/S) to terminate the enzymatic reaction. The mixture was centrifuged at 300 × *g* for 5 minutes, and the supernatant was discarded. The resulting pellet was resuspended in fresh DMEM containing 20% FBS and 1% P/S and seeded onto tissue culture plates.

Cells were incubated at 37°C in a humidified 5% CO₂ atmosphere and expanded until passage 5 (P5). From passage 1 (P1) onward, cultures were maintained in DMEM supplemented with 10% FBS but without antibiotics, to avoid potential effects of antimicrobial agents on cell proliferation or differentiation. All procedures were conducted under aseptic conditions using biosafety level 2 (BSL-2) practices.

### Soft Agar Colony Formation Assay

To evaluate anchorage-independent growth and confirm the non-transformed characteristics of the cells, a soft agar colony formation assay was performed using the CytoSelect™ 96-Well Cell Transformation Assay Kit (Cell Biolabs, Inc., San Diego, CA, USA), according to the manufacturer’s protocol.

HeLa, DPC, and NIS-DPC cells were cultured under standard conditions until reaching logarithmic growth phase. Cells were then harvested and resuspended in assay medium at the following densities: 8 × 10³ cells/well for HeLa, and 8 × 10⁴ cells/well for DPC and NIS-DPC. Cell suspensions were seeded in duplicate 96-well soft agar plates (Plate ① and Plate ②) as described in the kit instructions.

Plate ① was incubated for 7 days at 37°C in a humidified 5% CO₂ incubator to allow potential colony formation. Plate ② was processed immediately after cell seeding for background measurement following the manufacturer’s procedures: agar dissolution, cell lysis, and nuclear staining. The absorbance of each well was measured using a microplate reader to determine baseline optical density (OD).

After the 7-day incubation period, Plate ① was subjected to the same lysis and staining procedures, and absorbance was measured again. Colony formation was quantified by comparing OD values between the 0-day (Plate ②) and 7-day (Plate ①) samples. As expected, a significant increase in OD was observed only in the HeLa group, while DPC and NIS-DPC cells showed no increase, confirming their non-transformed phenotype.

### Construction of Lentiviral Vector pLVSIN-EF1α-hNIS-Pur

A self-inactivating (SIN) lentiviral expression vector, pLVSIN-EF1α-hNIS-Pur, was constructed based on the parental plasmid pLVSIN-EF1α-Pur (Takara Bio Inc.). The human sodium/iodide symporter (hNIS) gene (sequence ID No. 2; NCBI accession number NM_000453.3) was inserted into the multiple cloning site of the parental vector.

The DNA fragment encoding hNIS was synthesized with a Not I site at the 5′ end and a BamH I site at the 3′ end. Both the hNIS insert and the pLVSIN-EF1α-Pur vector were digested with the respective restriction enzymes Not I and BamH I (New England Biolabs or equivalent) to generate compatible cohesive ends. The digested fragments were ligated using Ligation High Ver. 2 (TOYOBO Co., Ltd.), yielding the recombinant plasmid pLVSIN-EF1α-hNIS-Pur.

All recombinant DNA manipulations were carried out under standard molecular cloning conditions in accordance with institutional biosafety regulations.

### Lentiviral Packaging and Production

Lentiviral particles carrying the pLVSIN-EF1α-hNIS-Pur construct were produced using a self-inactivating lentiviral packaging system. In addition to the expression vector described above, a mixture of packaging plasmids encoding HIV-1–derived structural and regulatory proteins (Gag, Pol, Tat, and Rev) and the vesicular stomatitis virus G glycoprotein (VSV-G) envelope was provided by the Lentiviral High Titer Packaging Mix (Takara Bio Inc.). Viral particles were generated following a modified standard protocol.

Poly-L-lysine–coated culture dishes were seeded with cells at a density of approximately 5 × 10⁴ cells/cm² and incubated overnight at 37°C in a humidified 5% CO₂ incubator. On the following day, a DNA mixture was prepared by adding 7 μL of Packaging Mix and 11 μL of pLVSIN-EF1α-hNIS-Pur plasmid DNA (0.5 μg/μL) to 1.5 mL of Opti-MEM I Reduced Serum Medium (Gibco).

Transfection was performed using either FuGENE HD (Promega) or TransIT-Lenti Transfection Reagent (Takara Bio Inc.), according to the manufacturers’ instructions. The transfected cells were incubated at 37°C in 5% CO₂ for approximately 24 hours, after which the culture supernatant was replaced with 10 mL of DMEM (Gibco) and incubation continued for another 24 hours.

The culture supernatant containing lentiviral particles was collected and clarified by centrifugation at 500 × *g* for 10 minutes. The supernatant was then passed through a 0.45 μm filter (Millipore, Cat. No. SLHPR33RB) to remove cellular debris. The filtered viral solution was aliquoted into microtubes and stored at −80°C until use.

### Generation of hNIS-Expressing Dental Pulp Cells (hNIS-DPCs)

DPCs were seeded at a density of 2.0 × 10⁴ cells/cm² and cultured overnight at 37°C in a humidified 5% CO₂ incubator. On the following day, cells were transduced with the hNIS lentiviral vector (hNIS-LV) at a multiplicity of infection (MOI) of 10. To enhance viral transduction efficiency, protamine sulfate (Sigma-Aldrich, Cat. No. P4020) was added to the viral inoculum at a final concentration of 100 μg/mL.

The viral supernatant was removed 24 hours after infection and replaced with fresh DMEM (Gibco, Cat. No. 10567-014). Cells were cultured with medium changes every 2–3 days, and cell morphology and confluence were monitored microscopically. When cultures reached approximately 90% confluence, cells were passaged using standard trypsinization procedures.

For selection of transduced cells, puromycin was added to the culture medium at a final concentration of 1 μg/mL during the first passage after viral infection. Resistant colonies were expanded under puromycin selection. Thereafter, cells were maintained in standard DMEM without selection antibiotic for one to two additional passages to establish a stable hNIS-DPC line. Established cells were cryopreserved in liquid nitrogen for long-term storage.

### Cell Culture

NIS-DPC and YTN16 cells were cultured in Dulbecco’s modified Eagle’s medium (DMEM; Gibco, Cat. No. 10567014) under standard conditions (37°C, 5% CO₂). Cells were maintained for 3 to 4 days before use.

### Cell Harvesting and Co-culture Setup

Cells were detached using 0.25% trypsin–EDTA (Gibco, Cat. No. 25200056) and resuspended in fresh DMEM. Cell concentrations were adjusted to 1 × 10⁵ cells/mL. For co-culture experiments, cells were seeded into 24-well Transwell plates (Corning, Cat. No. 353504) equipped with cell culture inserts (Corning, Cat. No. 351152). NIS-DPC cells (1 × 10⁵ cells) were seeded in the upper insert, and YTN16 cells (1 × 10⁵ cells) were seeded in the lower chamber. The co-cultures were incubated for 48 hours at 37°C in a humidified incubator with 5% CO₂.

### Migration Assay

After 48 hours of co-culture, the medium was aspirated and replaced with DMEM containing 1% human serum albumin (HSA) prepared from 25% clinical-grade albumin solution (Takeda Pharmaceutical Co., Ltd.). Cells were stained with Calcein (Dojindo, Cat. No. 341-07901) according to the manufacturer’s instructions to visualize viable cells. Fluorescence intensity was quantified using a spectrophotometric plate reader, and fluorescent images were captured using a fluorescence microscope under identical exposure conditions for all samples.

### Preparation of Mouse Model and Application of NIS-DPC Followed by Na[At-211]

Six-week-old female C57BL/6J mice were purchased from Jackson Laboratoy Japan and housed in a specific pathogen-free environment at the Animal Experiment Facility in Isotope Science Center, the University of Tokyo. The animal use proposal and experimental protocol were reviewed and approved by the University of Tokyo Animal Committee (ID: Med-P22-084、A2024ISC003). The YTN16 cell line was maintained in Dulbecco’s modified Eagle’s medium (DMEM, Sigma-Aldrich Merck, Tokyo, Japan) with 10% heat-inactivated fetal bovine serum (FBS, BWT, France), 50 μg/mL streptomycin, 50 U/mL penicillin (Thermo Fisher Scientific, Tokyo, Japan), and MITO + serum extender (Corning, NY, USA). A total of 1.0 ×10^7^ YTN16 cells were suspended in 500 μL of Hanks’ Balanced Salt Solution and intraperitoneally injected into mice (Ref. Nagaoka, 2025). Starting the following day, 2 μg of thyroxine was administered orally once daily. At 8, 10, and 12 days after YTN16 cell injection, 0.5 ×10^6^ NIS-DPC cells, 0.5 ×10^6^ DPC cells, or vehicle alone were administered intraperitoneally. On day 14, Na[At-211] (0.6 MBq) dissolved in 0.5 mL of vehicle was injected intraperitoneally. Nine days after the last Na[At-211] injection, the peritoneal cavity was examined, or the mice were observed until death or humane sacrifice, followed by examination of the peritoneal cavity.

### Cell Labeling with DiR and *In Vivo* Administration

NIS-DPC cells were suspended in medium at a concentration of 1 × 10⁷ cells/mL. The cell suspension was mixed with an equal volume of 20 μmol/L DiR dye solution (Thermo Fisher Scientific or equivalent), resulting in a final dye concentration of 10 μmol/L. The mixture was incubated for 15 minutes at room temperature in the dark, with gentle tapping every 5 minutes to ensure homogeneous labeling. After incubation, the cells were washed three times with injection medium by centrifugation to remove excess dye. The labeled cells were resuspended in injection medium at a final concentration of 1 × 10⁶ cells/mL. Labeled NIS-DPC cells were administered intraperitoneally to peritoneal dissemination model mice. All animal experiments were conducted in accordance with institutional guidelines and approved by the relevant Animal Care and Use Committee.

### *In Vivo* Fluorescence Imaging

At predetermined time points after injection, mice were sacrificed, and peritoneal tissues were harvested. DiR fluorescence signals were detected using an *in vivo* imaging system (IVIS Lumina III; PerkinElmer) under the following parameters: excitation filter, 740 nm; emission filter, 790 nm; and exposure time, automatic setting. Fluorescence images were analyzed using the manufacturer’s software.

### Production of At-211

According to the procedure described in Refs. (Yin, 2023, Yin, 2024), the radioisotope of At-211 was produced in the nuclear reaction of ^209^Bi(*α*,2*n*)^211^At using the AVF cyclotron of the RIKEN RI Beam Factory.

### Preparation of Na[At-211]

Dry solid At-211 of multiple chemical species in glass vial was dissolved the injection water (100 μL). At-211 aqueous solution (63 MBq, 50 μL) was mixed with 1% ascorbic acid, 2.3% MEYLON Injection aqueous solution (1 mL). After stirreing briefly, the reaction solution of At-211 was left to stand for 1hour at room temperature under sealed condition to afford the Na[At-211]. The Na[At-211] solution was dissolved 1% ascorbic acid, 2.3% MEYLON Injection aqueous solution, and used animal experiments and the cell tracer experiments.

### Thin-Layer Chromatography (TLC) Analysis

Thin-layer chromatography (TLC) was performed using silica gel 60 plates (Merck) as the stationary phase. Samples (5 μL, 280 kBq) were spotted onto the plate and developed in a solvent system consisting of acetonitrile and Milli-Q water (CH₃CN:MQ = 2:1, v/v).

After development, the chromatograms were analyzed using a TLC scanner (Scan-RAM; LabLogic Systems Ltd.) equipped with a ZnS scintillation detector. Radioactivity on the TLC plates was quantified using the manufacturer’s analysis software.

### Quantitative Measurement of At-211 Uptake

Na[At-211] (10 kBq) was added to the DPC and NIS-DPC cell and incubated for 5 minutes, 30 minutes, and 1 hour, respectively. After incubation, the medium was removed and the cells were collected. The radioactivity of the samples was then measured with a gamma-counter, and the counts per minute (CPM) were calculated.

### Visualization of alpha-ray

Alpha-particle track images were acquired with a CMOS sensor at 1-second exposure, yielding 600 frames each for NIS-DPS and DPC cells. Image analysis was conducted in ImageJ (v1.54f). For each frame, a fixed intensity threshold was applied, followed by morphological operations to generate a binarized mask of suprathreshold regions. Each image was then accumulated with its corresponding mask to isolate regions of interest surrounding the alpha-particle tracks, and the resulting stacks were Z-projected to produce composite track images.

## ACKNOWLEDGEMENTS

The radioisotope of At-211 was produced at the RI Beam Factory operated by the RIKEN Nishina Center and the CNS, University of Tokyo, Japan. The authors gratefully acknowledge Yousuke Kanayama, Nozomi Sato, Akihiro Nambu, Tomohiro Tomitsuka, Hiromichi Shimizu, Yudai Shigekawa, Sayantani Mitra, and Keiko Watanabe of RIKEN Nishina Center for technical assistance with the At-211 production.

## AUTHOR CONTRIBUTIONS

S.N. conceived and supervised the study and performed animal experiments. K.K., H.Y., H.M., T.Y., and W.-Y.D. conducted animal experiments. Y.W. performed alpha-particle therapy experiments and contributed to manuscript preparation. T.T. performed sodium At-211 synthesis. M.K. conducted cell culture experiments. A.T. performed RNA-seq data acquisition. T.Ta. analyzed pathological specimens. J.J. conducted At-211 compound analyses. Y.K. performed imaging studies. A.S. carried out alpha-particle uptake assays. R.N. prepared DPC cells and performed genetic modification of DPCs. W.Y. and H.S. performed functional analyses of DPCs and NIS-DPCs. K.I. managed DPC manufacturing and quality control. H.H. and X.Y. developed At-211 production technology. M.Yo. performed direct alpha-particle imaging. A.Yo. developed scintillators. T.K. and N.T. performed bioinformatics analyses. M.N. supervised bioinformatics and data integration. All authors discussed the results and approved the final manuscript.

## DATA AND MATERIALS AVAILABILITY

under preparation

